# NELLY enables patient-centric drug prioritization through interpretable drug-conditioned gene weighting

**DOI:** 10.64898/2026.08.25.747034

**Authors:** Christian Peralta Viteri, Nadja Harnischfeger, Lili Szabó, Stefan Hartmann, Kai Kretzschmar

## Abstract

Precision oncology seeks to match each tumor with the most effective anti-cancer therapy. Advances in pharmacogenomics and machine learning enabled drug response prediction models with strong performance in cancer cell lines. Nonetheless, patient-centric evaluation of drug prioritization and systematic assessment of model generalization in patient-derived systems across cancer types remain largely absent. Here we introduce a translational framework combining patient-centric benchmarking with a pan-cancer pharmacogenomic atlas of patient-derived organoids, together with NELLY, a deep learning model integrating transcriptomic and chemical information to predict drug response and prioritize therapies. NELLY outperformed existing methods for patient-specific drug prioritization across cancer cell lines and patient-derived organoids, including under out-of-distribution evaluation. Its dynamic weighting mechanism provided patient-specific gene attributions, offering a route to connect predicted drug response to molecular programs associated with drug resistance. Our results support NELLY as a promising framework for translationally relevant and interpretable drug response prediction in precision oncology.

---

Cancer remains a leading cause of death worldwide and poses a major challenge for treatment selection. Advances in oncology have highlighted the molecular heterogeneity that exists both across cancer types [1] and across patients with the same disease [2, 3]. These developments have driven the rise of precision/personalized medicine, which aims to use patient-specific molecular features to guide therapeutic decisions. In current clinical practice, treatment selection often relies on predictive biomarkers [4, 5, 6, 7]. However, this strategy has important limitations, patients may lack an actionable biomarker and, even when such biomarkers are present, clinical response may remain heterogeneous [8]. These challenges have motivated growing interest in predictive oncology models that use molecular profiling to anticipate treatment response [9, 10].

In this context, machine learning has emerged as a promising approach for the prediction of patients’ drug response. Accurate drug response prediction (DRP) from patient molecular data is a vital goal for precision medicine. Over the past decade, several machine learning methods have been developed for this task, with recent advances increasingly relying on deep learning for multimodal data integration. For example, DeepCDR [11] integrates genomics, transcriptomics, epigenomics and chemical structure through a hybrid graph-based architecture, whereas DualGCN [12] combines transcriptomics, copy number variation data and chemical structure using a dual graph convolutional framework. In parallel, other approaches have emphasized interpretability, particularly through attention-based architectures. For instance, Paccmann [13, 14] was introduced as a multi-head attention model integrating transcriptomics, chemical structure and prior knowledge network. Nonetheless, Jin et al. (2021) [15] argued that attention scores in Paccmann were not always biologically consistent across related genes, thereby limiting biological interpretation. To address this, they proposed HiDRA, a hierarchical attention network that represents biological pathways and the genes within them related to drug response.

Despite this progress, several studies have highlighted key limitations that hinder their application to precision oncology. Firstly, many methods have not been evaluated under sufficiently robust experimental designs. This concerns both the data-splitting strategy and the performance metrics in the evaluation framework. In particular, unstratified evaluations have led to inflated performance estimates [16, 17]. As a result, patient-centric evaluation of model performance remains scarce, despite its importance for precision oncology [18, 19].

A second major limitation is the lack of systematic evaluation of DRP models on patient-derived data across cancer types [17, 20]. Although previous studies have reported promising performance across cancer cell line (CCL) datasets [17, 21], the application of DRP models for translational purposes is still limited. Personalized decision-making requires evaluation on cancer models that recapitulate patient-specific features more faithfully. In this context, patient-derived organoid (PDO) technology has established itself as a powerful platform to study personalized anti-cancer therapy [22, 23, 24]. PDO lines can reproduce patient-specific tumor heterogeneity [25, 26] and predict patient response to therapy [27, 28]. Although previous work has tested their predictive model on PDO lines from a specific cancer type [19, 29], there is no comprehensive resource available for the systematic evaluation of DRP generalization across cancer types and therapeutic agents.

Lastly, interpretability is a key requirement for precision oncology, as it enables both interpretation of model predictions and biologically informed hypothesis generation [30]. Attention-based architectures have gained popularity in DRP, due to their expressive power and potential to provide gene-level information. However, despite improvements in biological structure and interpretability, such as those introduced in HiDRA [15], current approaches offer limited insight into which molecular features drive an individual patient’s predicted response. Instead, their representations often have to be interpreted as a cohort-level summary. This highlights the need for methods that enable gene attribution to be explored at the patient-specific level.

Here, we adopted recent best practices for robust DRP evaluation [16, 19] and, in addition, implemented rank-based metrics for drug recommendation to address these limitations. Furthermore, we assembled a pan-cancer PDO-based pharmacogenomic atlas for out-of-distribution (OOD) generalization assessment and developed a new neural network module that enables patient-specific gene attribution investigation.

## Results

### Translationally relevant benchmark for drug response prediction

Previous work had shown that gene expression, compared to other “-omic” modalities, provides superior predictive power for this task [31]. Motivated by these results, we selected transcriptomic data as the molecular features for our precision-medicine-oriented drug response prediction framework.

To mimic a more clinically-relevant setting, the evaluation of models on unseen-CCL was recently proposed [29]. In this framework, training data were split using an unseen-CCL 10-fold cross-validation scheme, in which models were fitted on a set of CCLs and evaluated on previously unseen-CCLs (Fig. 1A, top). In addition to Pearson correlation as a measure of prediction accuracy, ranking-based metrics were included to quantify how accurately models produce drug recommendations (Fig. 1A, bottom). We reasoned that combining regression and recommendation metrics would provide complementary perspectives for assessing the extent to which DRP models can support precision oncology.

**Figure 1:**
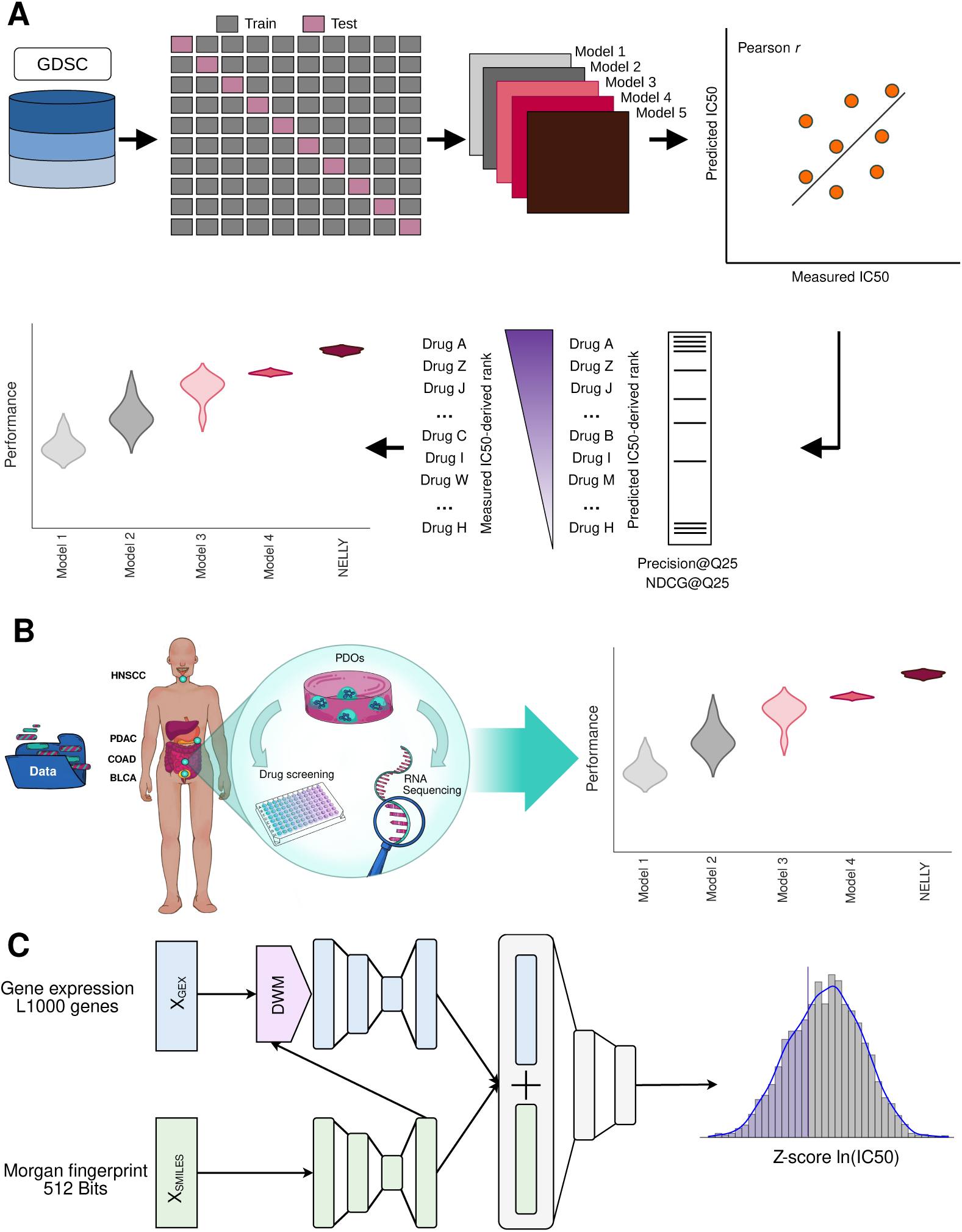
Translationally relevant benchmark for drug response prediction. **A.** Study design applied for patient-centric model training and evaluation. Model performance was compared both in terms of regression accuracy (Pearson correlation coefficient) and ranking accuracy (Precision@Q25 and NDCG@Q25). **B.** Assembly of a pan-cancer patient-derived organoid (PDO) dataset. Gene expression and drug screening data were collected from publicly available datasets, or generated in this study. This PDO-based pharmacogenomic atlas was used as an independent validation to benchmark model performance under out-of-distribution evaluation. **C.** Neural network architecture. NELLY consisted of a multimodal neural network with three dedicated heads. The first two heads extracted informative patterns from gene expression (GEX) and SMILES and projected them into an embedding space. GEX and SMILES embeddings were then summed and fed into the third head, the prediction head, to predict a Z-score ln(IC50) value.

A translationally relevant evaluation framework is essential for personalized medicine. Although several studies have carried out drug screenings in PDOs, harmonized pharmacogenomic resources for PDO-based model evaluation remain largely absent. To address this gap, a pan-cancer PDO-based pharmacogenomic dataset was assembled from published studies with high-quality drug-screening data [25, 32, 33, 34] and this study. We then used this resource as an independent validation set to benchmark model performance under OOD evaluation (Fig. 1B).

We developed NELLY as a deep learning model for robust generalization and drug prioritization on PDO lines (Fig. 1C). NELLY’s architecture was designed as a multimodal neural network that integrated baseline gene expression data (GEX) with drug chemical features derived from SMILES representations. The GEX consisted of log-normalized expression values for the L1000 genes, a compact yet biologically informative gene set covering diverse biological processes [35]. In parallel, SMILES were encoded in the form of 512 bits Morgan fingerprints [36]. The architecture consisted of three dedicated heads. The first two heads encoded the GEX and SMILES, extracting meaningful patterns and mapping information to a latent space. GEX and SMILES embeddings were then summed and fed into the third head, the prediction head, to predict a Z-score ln(IC_50_) value for a given CCL-drug pair (Fig. 1C).

### DWM-NELLY outperforms existing models on unseen cancer cell lines

To benchmark NELLY under comparable and methodologically consistent conditions, we selected three existing deep learning models for drug response prediction: Paccmann [13, 14], HiDRA [15] and ScreenDL [29]. These models were chosen because they were developed using GDSC drug response data and share the same core input modalities as NELLY: gene expression and drug chemical information, thereby making them directly comparable. At the same time, they represent distinct state-of-the-art strategies for DRP, including multimodal multi-head attention-based integration, pathway-informed hierarchical attention and deep neural drug-response modeling. NELLY was designed to address a current limitation of these approaches: the need to couple accurate prediction with patient-level molecular interpretation. Rather than relying on latent or pathway-level representations that are commonly interpreted after aggregation across samples providing a cohort-level summary, NELLY uses a drug-conditioned gene-weighting mechanism that modulates transcriptomic features directly for each sample-drug pair. This preserved a direct correspondence between model-derived weights and input genes, enabling patient-level investigation of molecular features associated with predicted drug response. NELLY therefore extends drug response prediction by providing a fine-grained interpretability layer, addressing an important requirement for predictive precision oncology.

We introduced the Dynamic Weighting Mechanism (DWM) (Fig. 2A), that took as input the GEX matrix (*X*_GEX_) and the SMILES-derived chemical embedding (*Z*_SMILES_). These two matrices were concatenated and passed through a learned linear projection parameterized by a weight matrix (*W*), producing an attribution matrix (*S*). This matrix was then applied to *X*_GEX_ via element-wise multiplication, serving as a gating mechanism that up- or down-weighted genes based on their relevance to the prediction task. To improve training stability and preserve the original transcriptomic signal, the DWM output was summed with *X*_GEX_ through a residual connection [37].

**Figure 2:**
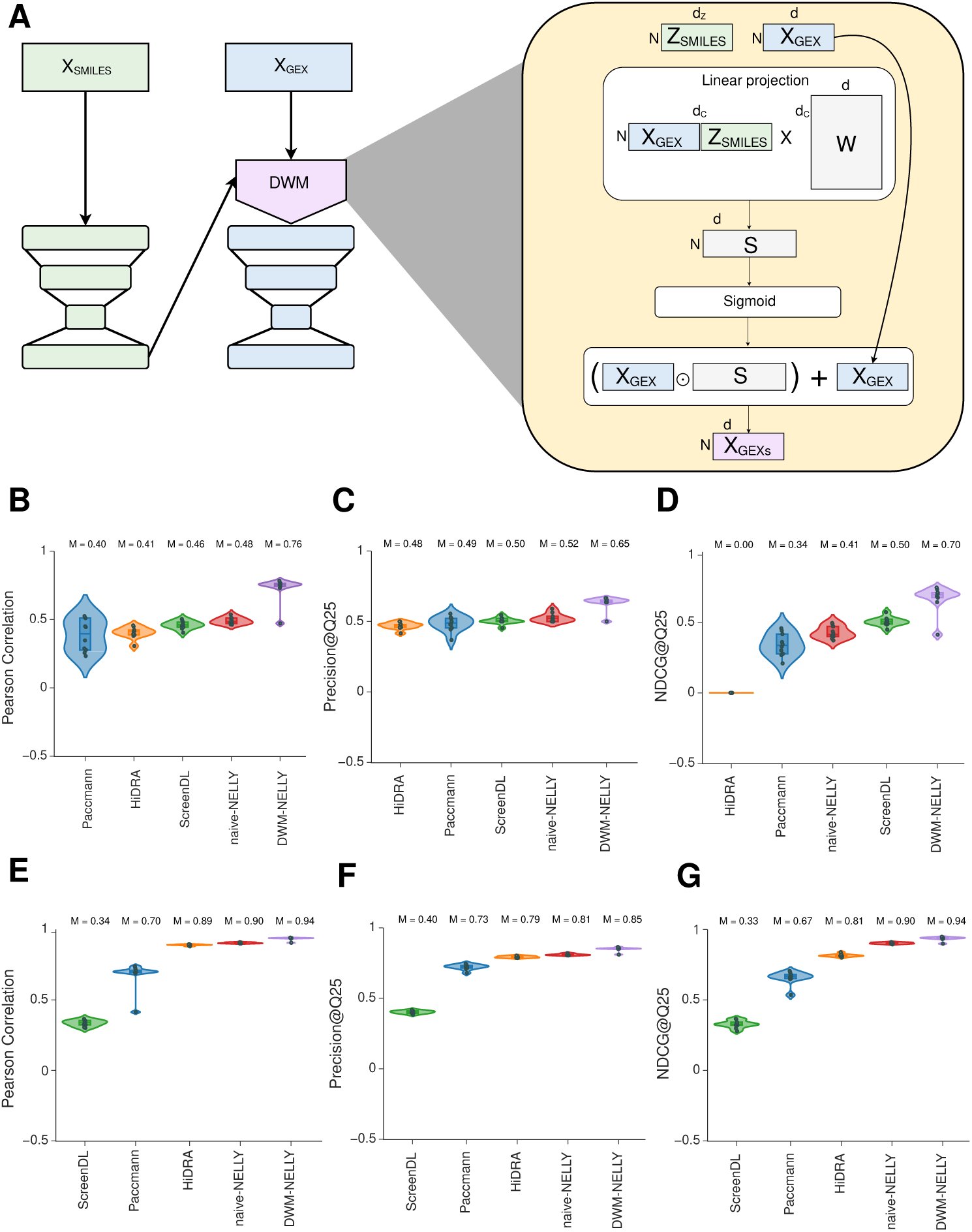
DWM-NELLY outperforms existing models on unseen cancer cell lines. **A.** Dynamic Weighting Mechanism (DWM) scheme. The DWM took as input the gene expression matrix (*X_GEX_*) and SMILES-derived chemical embedding (*Z_SMILES_*). These two matrices were concatenated and passed through a learned linear projection, producing a gene attribution matrix (*S*). The *S* matrix was then applied to *X_GEX_* via element-wise multiplication, modulating gene contribution to the prediction task. To improve training stability, the DWM output was combined with *X_GEX_* via a residual connection. **B-D.** Fixed-drug aggregation benchmark. Performance was evaluated both as regression accuracy (Pearson correlation coefficient) and ranking accuracy (Precision@Q25 and NDCG@Q25). This setting assessed the model’s ability to predict across different biological contexts for a given compound. This perspective quantified how well the model captured sensitivity patterns induced by the same drug. **E-G.** Fixed-lined aggregation benchmark. Performance was evaluated both as regression accuracy (Pearson correlation coefficient) and ranking accuracy (Precision@Q25 and NDCG@Q25). This setting assessed the model’s ability to predict across different drug perturbations within the same cancer line. This perspective quantified the model’s ability to capture cell line-specific drug responses.

The benchmark was performed on an unseen-CCL 10-fold CV scheme, as described in the previous section, and the Pearson correlation coefficient (PCC) was used to comply with standard evaluation metrics. To complement regression-based evaluation, model predictions were also assessed as ranked drug-response recommendations. Following the Q25 sensitivity criterion defined in the Methods, precision at Q25 (Precision@Q25) was used to quantify the recovery of the most sensitive responses, whereas Normalized Discounted Cumulative Gain at Q25 (NDCG@Q25) was used to evaluate the ordering of the predicted values. Each metric was aggregated according to two strategies, i.e, fixed-drug and fixed-line [16]. For model comparison, the median was computed per fold and statistical comparisons were performed using a two-sided Wilcoxon signed-rank test (Fig. 2B-G). This benchmark allowed us to evaluate whether the drug-conditioned gene-weighting strategy implemented in NELLY provided advantages over existing architectures.

On one hand, fixed-drug aggregation allowed us to assess the model’s ability to predict different drug perturbations between heterogeneous biological context for a given compound. This perspective emphasized how well the model captured sensitivity patterns induced by the same drug (Fig. 2B-D). Under this strategy, ScreenDL has been reported as the most accurate model, achieving a median PCC of 0.46 [29] (Fig. 2B). However, DWM-NELLY scored a median PCC of 0.76, a significant improvement (p-value = 1.9531 × 10*^−^*^3^, Wilcoxon signed-rank test). For the recommendation metrics, DWM-NELLY achieved median scores of 0.65 and 0.70 for Precision@Q25 and NDCG@Q25, respectively. This indicated that NELLY not only identified relevant CCL responses among the top-ranked predictions, but also ranked them more accurately, as measured by NDCG@Q25. These results indicated a significant improvement in NDCG@Q25 compared to ScreenDL (0.70 vs 0.50, p-value = 1.9531 × 10*^−^*^3^, Wilcoxon signed-rank test).

In contrast, fixed-line aggregation allowed us to assess the model’s ability to predict different drug perturbations within the same line. This perspective emphasized how well the model captured CCL-specific drug responses (Fig. 2E-G). DWM-NELLY showed a significant improvement with a median PCC = 0.94, compared to the best existing model, HiDRA (0.94 vs 0.89, p-value = 9.7656 × 10*^−^*^4^, Wilcoxon signed-rank test). For the recommendation metrics, DWM-NELLY achieved median scores of 0.85 and 0.95 for Precision@Q25 and NDCG@Q25, respectively. This indicated that NELLY not only identified the most sensitive drugs more effectively, but also ranked them more accurately. Consistently, DWM-NELLY outperformed HiDRA in NDCG@Q25 (0.94 vs 0.81, p-value = 1.9531 × 10*^−^*^3^, Wilcoxon signed-rank test).

### DWM-NELLY learns the pharmacological landscape and the contextual mechanism of action

To better understand what DWM-NELLY learned about drug response, we examined the *in-silico* pharmacological landscape. Specifically, all 413 drugs were represented as a vector of predicted Z-score ln(IC_50_) values across all 952 CCLs [38] (Fig. 3A). When this high-dimensional matrix was projected into a 2D representation using Uniform Manifold Approximation and Projection (UMAP) [39], drugs were organized along a continuous manifold. This primary axis of variation aligned with the average of the observed Z-score ln(IC_50_) of each drug, indicating that the model captured global trends in the pharmacological landscape (Fig. 3B). This organization suggested that DWM-NELLY learned a coherent space, distinguishing between broadly cytotoxic versus broadly selective compounds.

**Figure 3:**
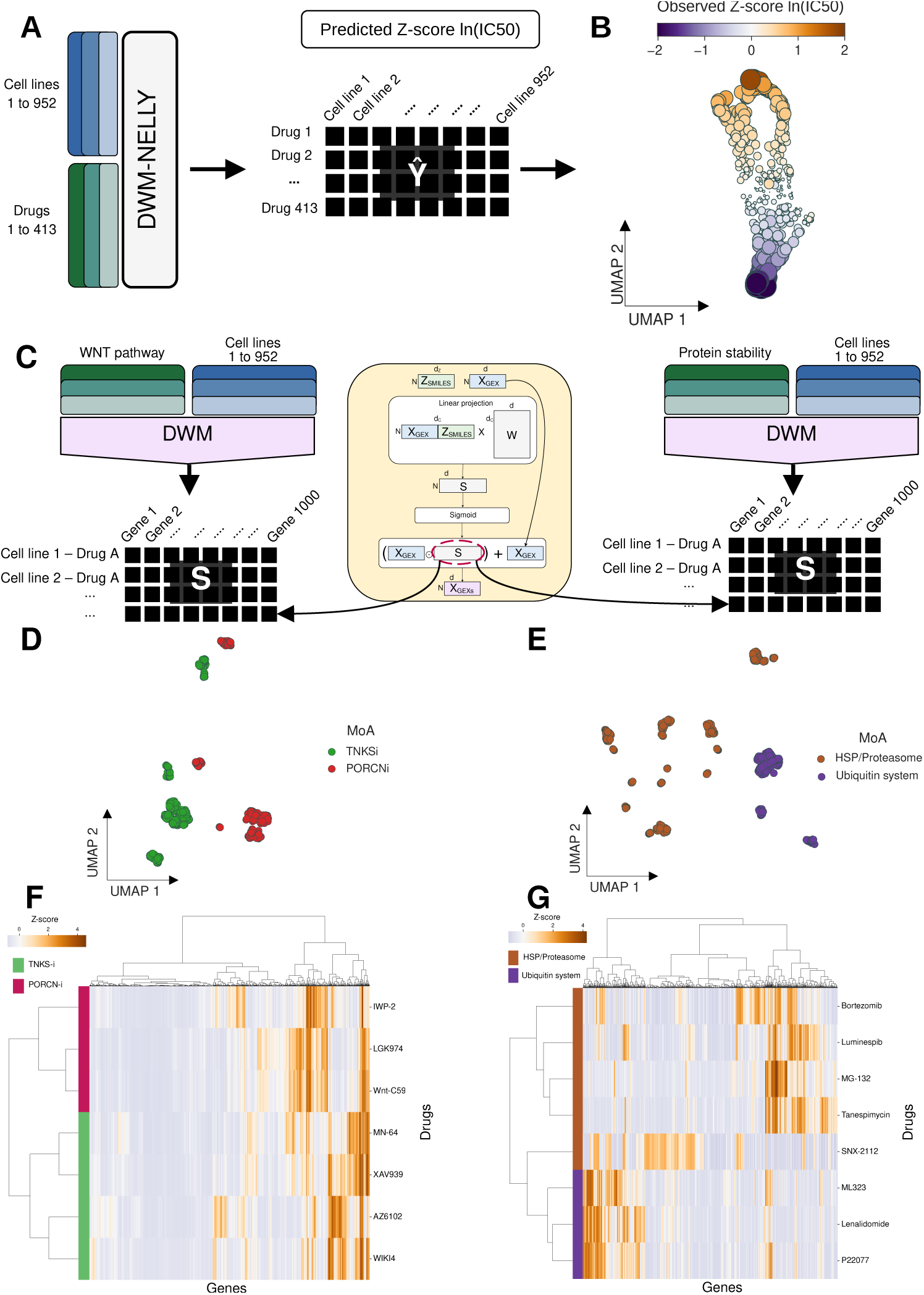
DWM-NELLY learns the pharmacological landscape and the contextual mechanism of action. **A.** Pre-trained NELLY was used to predict drug response for the training data. Predicted Z-score ln(IC50) values were arranged into a drug-by-cell-line matrix, where each drug was represented by a vector of bioactivity across 952 cell lines. **B.** Drug bioactivity space. The bioactivity matrix was projected using UMAP to obtain a 2D representation in which drugs were organized along a continuous manifold. The primary axis of variation aligned with the average observed Z-score ln(IC50) of each drug, indicating that the model captured global trends in the pharmacological landscape. **C.** DWM-derived gene attribution matrix (*S*) was extracted for drugs targeting the WNT pathway (left) and protein stability (right). **D.** WNT inhibitors space. The gene attribution matrix was projected using UMAP to obtain a 2D representation that shows separation by drug mechanism of action for WNT inhibitors. **E.** Protein stability inhibitors space. The gene attribution matrix was projected using UMAP to obtain a 2D representation that shows separation by drug mechanism of action for protein stability inhibitors. **F.** WNT inhibitors clustering. Hierarchical clustering of the drug-averaged attribution matrix further revealed drug groupings based on model-assigned gene weights for WNT inhibitors. **G.** Protein stability inhibitors clustering. Hierarchical clustering of the drug-averaged attribution matrix further revealed drug groupings based on model-assigned gene weights for protein stability inhibitors.

Next, we interrogated the role of the DWM in determining the drug mechanism of action (MoA). In particular, drugs were selected based on their target pathway classification, focusing on classes with multiple MoA. Accordingly, therapies targeting single proteins or broadly promiscuous signaling pathways were excluded. In total, seven drugs targeting WNT signaling and eight drugs targeting protein stability/degradation pathways were selected for analysis. The corresponding attribution matrices (*S*) were extracted from the DWM (Fig. 2A).

Here, each CCL-drug pair was represented as a vector of attributed weights across 1000 genes, separately for drugs targeting the WNT pathway (Fig. 3C, left), and protein stability/degradation (Fig. 3C, right). UMAP projection of these representations showed separation between the two MoA present within WNT-targeting drugs, i.e., TNKS inhibitors and PORCN inhibitors (Fig. 3D). Similarly, protein stability/degradation UMAP projection showed a separation between drugs inhibiting heat shock proteins (HSP) and proteasome, from those acting on the ubiquitin system (Fig 3E). Consistently, hierarchical clustering of the drug-averaged attribution matrix (*S*) revealed separation between drugs driven by model-attributed gene weights (Fig. 3F and 3G).

These results showed that DWM-NELLY learned biologically coherent representations of drug response, enabling the distinction between broadly sensitive and broadly resistant compounds. In addition, gene contributions generated by the DWM were consistent with drug MoA. Taken together, these findings suggested that transferable drug response representations had been learned, motivating the subsequent evaluation of DWM-NELLY under OOD prediction in PDO lines, which represented a distinct and more patient-relevant data distribution.

### DWM-NELLY generalizes to patient-specific IC_50_ prediction in PDOs

To evaluate whether DRP models generalize to patient-relevant material, we assembled a pan-cancer PDO-based pharmacogenomic dataset and used it as an independent benchmark for patient-specific IC_50_ prediction. The PDO benchmarking dataset included data from 19 PDO lines from colorectal adenocarcinoma (COAD) [25], 11 PDO lines from bladder cancer (BLCA) [32], 5 PDO lines from pancreatic ductal adenocarcinoma (PDAC) [33], 7 PDO lines from head and neck squamous cell carcinoma (HNSCC) [34] and 12 HNSCC PDO lines generated in this study (Fig. 4A). Drug response data included the IC_50_ values for 129 different drugs covering 28 different target pathways (Fig. 4B).

**Figure 4:**
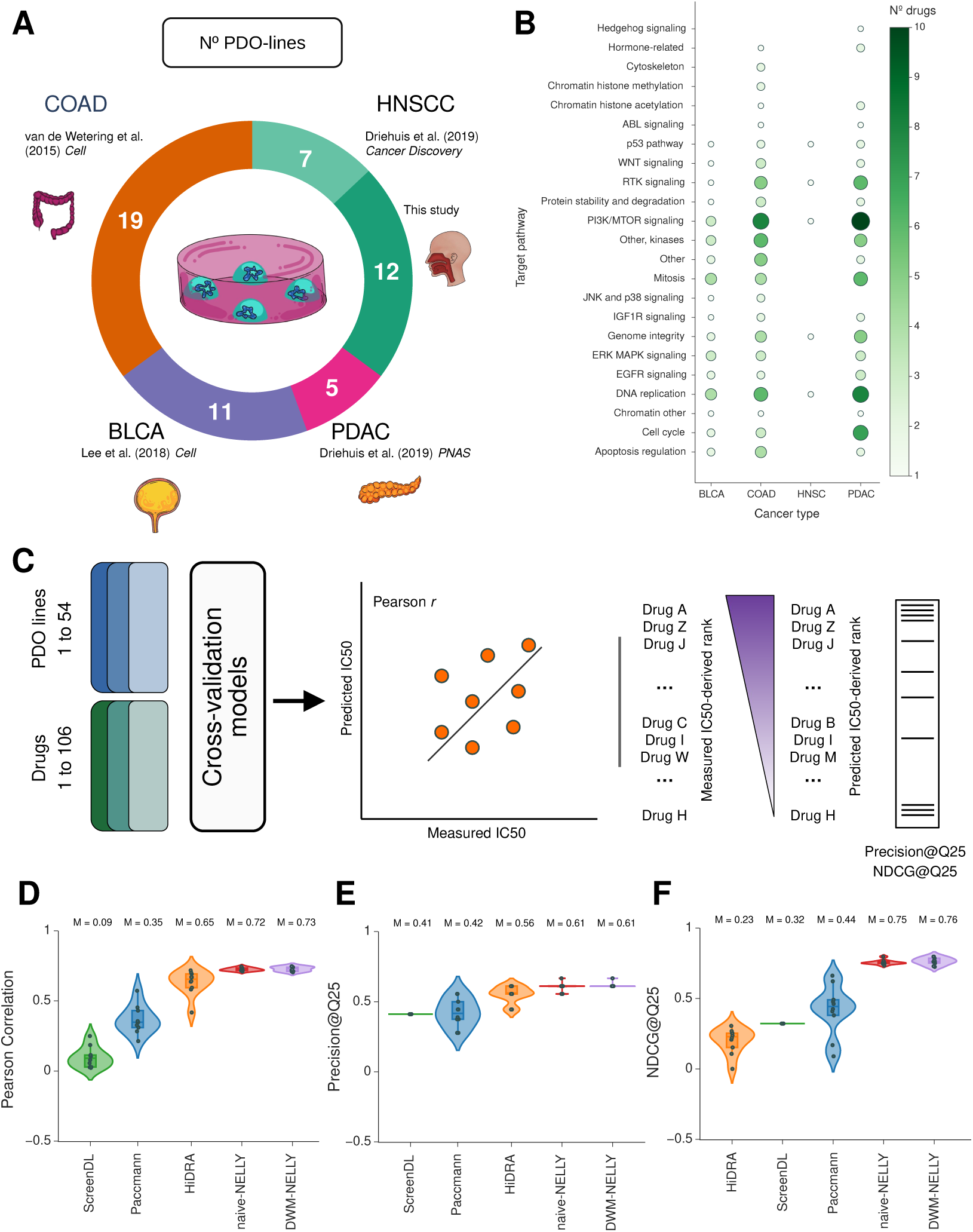
DWM-NELLY generalizes to patient-specific IC50 prediction in organoid lines. **A.** Overview of the pan-cancer patient-derived organoid pharmacogenomic atlas. The dataset contained organoid lines from 4 tumor entities and 5 studies. **B.** Overview of the therapeutic agents included in the atlas. The atlas contained compounds targeting 21 established pathways. **C.** Generalization benchmark scheme. Model performance was assessed under out-of-distribution prediction. Performance was evaluated both as regression accuracy (Pearson correlation coefficient) and ranking accuracy (Precision@Q25 and NDCG@Q25). **D-F.** Fixed-line aggregation benchmark on patient-derived organoid dataset. This patient-centric aggregation strategy quantified the accuracy with which models predict patient-specific drug responses and generate drug prioritization.

The generalization evaluation strategy was depicted in Figure 4C. Models obtained from the cross-validation were applied to PDO-drug pairs to generate drug response predictions under OOD evaluation. Generalization performance was assessed both as regression accuracy (the Pearson correlation coefficient) and as ranking performance (Precision@Q25 and NDCG@Q25).

For a precision oncology application, we conducted a model benchmark under fixed-line aggregation (Fig. 4D-F). This allowed us to assess the model’s ability to detect top sensitive drugs in a patient-specific manner. Under regression-based evaluation, HiDRA was the best-performing existing model, achieving a median PCC = 0.65 (Fig. 4D). Under ranking-based evaluation, Paccmann was the best-performing existing model, achieving a median NDCG@Q25 = 0.44 (Fig. 4F). Although existing models showed robust generalization to PDO lines, DWM-NELLY consistently achieved higher scores. In terms of regression performance, DWM-NELLY outperformed HiDRA (0.73 vs 0.65, p-value = 1.9531 × 10*^−^*^3^, Wilcoxon signed-rank test). Whereas in ranking performance, DWM-NELLY outperformed Paccmann (0.76 vs 0.44, p-value = 1.9531 × 10*^−^*^3^, Wilcoxon signed-rank test).

Taken together, these results demonstrated that DWM-NELLY is a robust framework for drug response prediction. DWM-NELLY achieved the highest scores across benchmarks, and the contextualization analysis in the previous figure further suggests that the model learned coherent drug-specific representations related to the mechanism of action. Its strong generalization to PDOs further supported its potential for patient-specific drug prioritization.

### DWM uncovers regulatory programs underlying therapeutic resistance

Despite advances in the field, HNSCC remains therapeutically challenging. To investigate whether our DWM can support the study of drug response beyond prediction alone, we extracted the DWM-derived gene attribution (*S*) (Fig. 2A). First, we performed *in-silico* drug screening, pairing each HNSCC PDO line with the 413 drugs used during training (Fig. 5A, left). To ensure robustness of the downstream analysis, PDO-drug pairs were fed into the 10 models obtained from the cross-validation procedure, obtaining the predicted Z-score ln(IC_50_). The resulting predictions were then aggregated by taking the median value across models for each PDO-drug pair. By aggregating across the 10 models, we approximated an ensemble prediction and reduced dependence on a single trained model.

**Figure 5:**
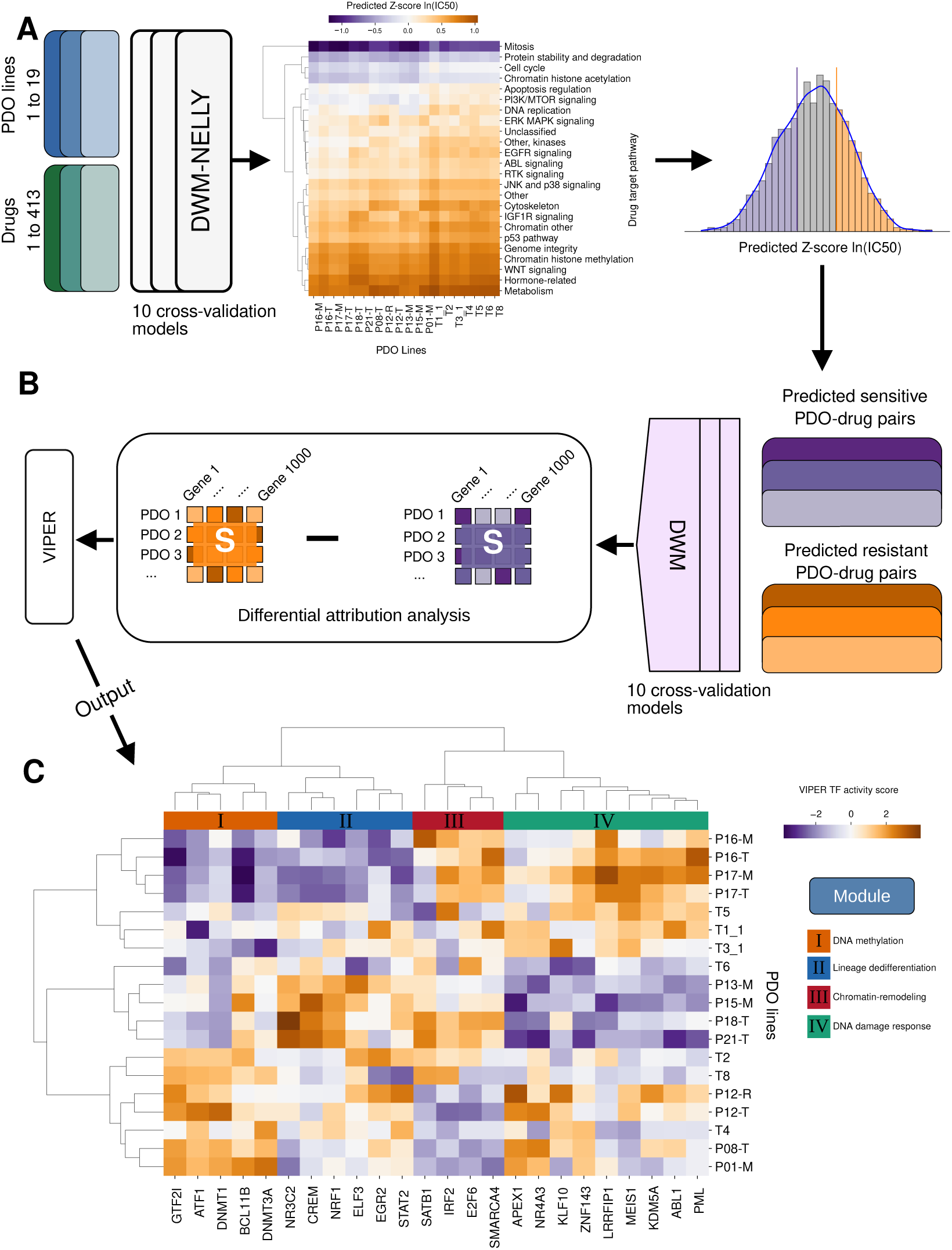
DWM uncovers regulatory programs underlying therapeutic resistance. **A.** *In-silico* drug screening on HNSCC cohort. Each HNSCC patient-derived organoid line was paired with every drug included in the training set to generate DWM-NELLY predictions. For each organoid line, we then extracted the drug responses within into the lower 25% and upper 75% of the predicted Z-score ln(IC50) distribution. **B.** Differential gene attribution analysis. Predicted sensitive and predicted resistant PDO-drug pairs were passed through the DWM to obtain gene attribution matrices. These attribution matrices were then compared to identify genes assigned higher weights to predicted-resistant than to predicted-sensitive responses. The resulting differential values were used to infer transcription factor activity using the VIPER algorithm. **C.** VIPER output. Heatmap of inferred transcription factor activity showing 4 resistance-related clusters.

Predicted drug responses recapitulated the well-described pattern of broad drug insensitivity in HNSCC (Fig. 5A, center) [40, 41, 42]. For each PDO line, predicted responses were stratified as sensitive or resistant when the predicted Z-score ln(IC_50_) value fell within the lower 25% or upper 75% of the distribution, respectively (Fig. 5A, right). Similarly to the predicted drug response, the predicted-sensitive and predicted-resistant PDO-drug pairs were passed through the DWM of the 10 models obtained from the cross-validation (Fig. 5B, right). Each *S* matrix was then aggregated by taking the median gene contribution values across models for each PDO-drug pair. The *S* matrices were then used to identify genes assigned higher weights in predicted-resistant responses than in predicted-sensitive responses (Fig. 5B, center). Finally, these differential attribution values were used to infer transcription factor (TF) activity using the VIPER algorithm [43] (Fig. 5B, left). VIPER estimates regulator activity from the coordinated behavior of downstream target genes rather than from the expression of the transcription factors.

Hierarchical clustering analysis of TF activity revealed four major regulatory modules, indicating that resistance was not driven by a single factor, but by a combination of cellular programs (Fig. 5C).

Cluster I (*GTF2I, ATF1, DNMT1, BCL11B, DNMT3A*) was characterized as a DNA methylation-dependent module. *DNMT1* and *DNMT3A* encode DNA methyltransferases primarily involved in maintenance and *de novo* methylation, respectively [44, 45]. In HNSCC, aberrant DNA methylation has been repeatedly implicated in therapy adaptation [46, 47, 48]. Decitabine-mediated inhibition of DNA methylation has been reported to restore Cisplatin sensitivity in resistant models [49]. Accordingly, cluster I was interpreted as an epigenetic stabilization module. Cluster II (*NR3C2, CREM, NRF1, ELF3, EGR2, STAT2*) showed an intermediate pattern, with fewer well-established resistance mediators (Fig. 5C). *ELF3* is an epithelial lineage-associated TF [50], whereas *STAT2* is a core component of type I interferon signaling [51]. Together, these factors suggest that this module might reflect altered epithelial differentiation and JAK/STAT-related signalling which could constitute a putative axis of cell-death evasion.

Cluster III (*SATB1, IRF2, E2F6, SMARCA4*) seemed to describe a chromatin-remodeling module (Fig. 5C). In contrast to cluster I, this module was not centered on methylation-dependent regulations. Instead, it was characterized by *SATB1*, encoding a chromatin organizer [52, 53], and *SMARCA4*, enconding the ATPase subunit of the SWI/SNF chromatin-remodelling complex [54]. *SATB1* has been associated with aggressive squamous phenotypes and poor prognosis, while *SMARCA4* has been linked to invasion and drug-tolerant states [55, 56, 57].

Cluster IV (*APEX1, NR4A3, KLF10, ZNF143, LRRFIP1, MEIS1, KDM5A, ABL1, PML*) also comprised several well-described resistance-associated candidates (Fig. 5C). In particular, *APEX1* plays a central role in DNA repair and has been shown to be relevant to the DNA damage response in HNSCC [58, 59]. *ZNF143* has mechanistic links to Cisplatin resistance and the DNA damage-response [60]. *KDM5A* encondes a mediator of reversible drug-tolerant states across solid tumors [61, 62, 63, 64], and *ABL1* has been implicated in radiation resistance in HNSCC [65].

Overall, our analysis of TF activity from predicted resistant samples supported a model in which drug resistance might be induced by at least two complementary layers. The first layer consists of epigenetic and chromatin-regulatory programs, represented by cluster I and cluster III, with distinct underlying mechanisms: DNA methylation in cluster I and chromatin remodeling in cluster III. The second layer was more closely related to repair-associated and stress-adaptive programs, represented by cluster IV. Cluster II appeared intermediate and was less readily interpreted as a canonical resistance module, but may reflect altered epithelial and interferon-responsive transcriptional states accompanying resistance. Together, these results suggested that DWM-derived gene attribution might provide mechanistic interpretability beyond drug ranking by resolving biologically coherent regulatory modules associated with therapeutic resistance in HNSCC.

### DWM-NELLY predicts patient-specific drug recommendations in HNSCC

To investigate whether DWM-NELLY could support personalized drug prioritization for HNSCC patients, we leveraged the *in-silico* drug screening described in the previous section. To detect personalized drug recommendations, we selected drugs whose predicted Z-score ln(IC_50_) values fell within the lower 25% of the distribution for each PDO line (Fig. 5A, right). Next, we computed a normalized rank score within the recommended drug set, linearly scaling ranks to the [0, 1] interval, where 1 denotes the top-ranked drug and 0 the lowest-ranked drug within Q25 for each PDO line independently (Fig. 6A). This strategy was applied to focus on therapeutic agents that can constitute promising candidates to elicit a sensitive response in individual PDO lines.

**Figure 6:**
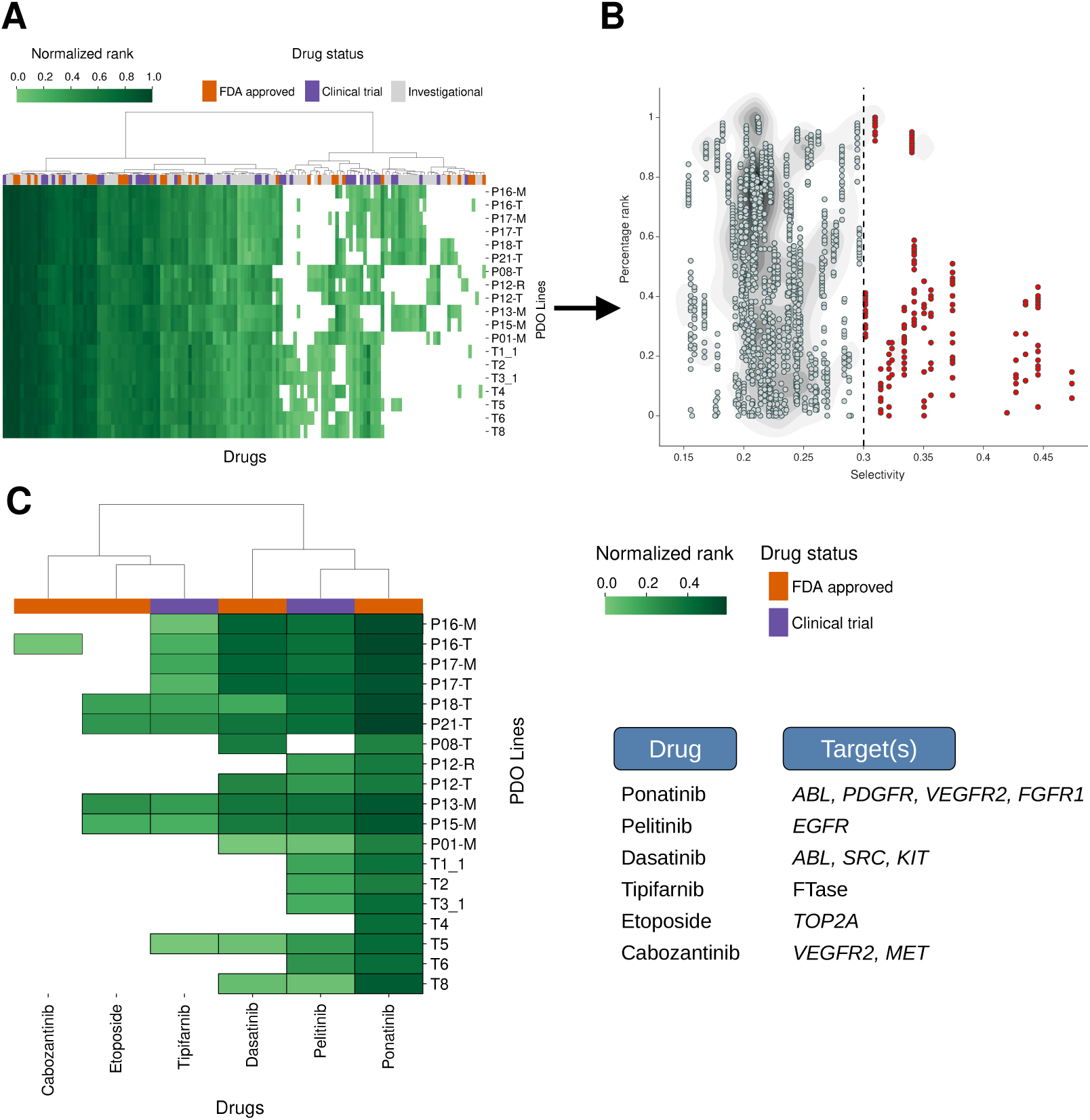
DWM-NELLY predict patient-specific drug recommendations in HNSCC. **A.** Drug response ranking. Predicted drug responses were transformed into a normalized rank score, from 0 to 1. Values closer to 1 indicate higher predicted sensitivity, whereas values closer to 0 indicate lower predicted sensitivity within the selected lower 25%. **B.** Drug selectivity assessment. Drug recommendations were further filtered to distinguish broad cytotoxicity from selective activity. Using the bimodality coefficient as a selectivity criterion enabled the prioritization of drugs with potential selective anti-cancer effect for the HNSCC cohort. **C.** Drug prioritization. Final patient-specific drug recommendations generated by the DWM-NELLY framework.

A critical challenge in DRP for precision oncology is distinguishing drugs with broad cyto-toxic activity from those with a more selective activity profile [66, 67]. This distinction is important for prioritizing compounds with greater patient specificity. To address this, we applied the selectivity criterion introduced by Corselo et al. (2020) [68]. This criterion was based on the bimodality coefficient [69], which quantifies the tendency of a distribution to deviate from unimodality. Here, it was computed for each drug across PDO lines using the predicted Z-score ln(IC_50_) values. By combining this selectivity metric with normalized rank score, we aimed to distinguish drugs which were more likely to display broadly cytotoxic activity from drugs that were more likely to exhibit selective activity towards HNSCC PDO lines (Fig. 6B). In this context, compounds with more selective activity were expected to better exploit patient-specific vulnerabilities.

DWM-NELLY prioritized several therapeutic agents with prior biological or clinical relevance in HNSCC, supporting the biological plausibility of its recommendations (Fig. 6C). Among them, Pelitinib, an EGFR inhibitor, was in line with previous studies indicating an overexpression in 80% of cases which made it a suitable candidate for a phase 1 clinical trial [70, 71, 72]. However, clinical benefit from EGFR-targeted treatment has been variable. DWM-NELLY also suggested Cabozantinib, which inhibits VEGFR2 and MET, proteins associated with tumor aggressiveness and resistance to EGFR inhibitors [73, 74, 75]. Dasatinib, which inhibits SRC family kinases and ABL, has also attracted interest in HNSCC. *SRC* overexpression has been reported in HNSCC cell lines, and inhibition increased radiation sensitivity [76]. In addition, DWM-NELLY prioritized Etoposide, a topoisomerase inhibitor previously explored in a phase 1 clinical trial [77] and discussed in recent studies [41]. Lastly, the prioritization of Tipifarnib was particularly notable. HRAS inhibition has been the subject of multiple studies and one clinical trial for the precision treatment of a subset of HNSCC patients [7, 78, 79, 80]. However, a recent publication showed that Tipifarnib sensitivity might not be solely associated with HRAS mutation in PDO lines [81].

Taken together, these results indicated that DWM-NELLY prioritized biologically grounded therapeutic candidates in a personalized context, including compounds targeting established HNSCC-relevant pathways and previously explored clinically. This supports DWM-NELLY’s biological and translational relevance for prioritizing drugs despite the broadly insensitive pharmacological landscape of HNSCC.

## Discussion

In this study, we established a patient-centric framework for benchmaking DRP models and applied it in a translationally relevant setting using a pan-cancer PDO-based pharmacogenomic atlas. This framework provided a more informative view of model performance than conventional unstratified evaluation by emphasizing patient-specific drug prioritization and independent validation on patient-derived material. Under this setting, NELLY substantially outperformed existing methods [13, 15, 29] in patient-specific drug prioritization across both CCLs and PDOs. In addition, the DWM introduced an interpretability layer that enabled patient-specific gene attribution extraction and hypothesis generation beyond prediction alone.

Patient-centric evaluation is particularly important for translational application, as the clinical objective is not simply to predict a drug response, but to identify and rank the most suitable therapeutic options for an individual patient [16, 19]. For this reason, evaluation strategies for precision oncology should emphasize ranking accuracy in addition to regression performance. Similarly, assessing models on patient-derived systems is essential to determine whether predictive signals learned from preclinical data transfer to more clinically relevant cases. Together, these two strategies provide a clearer estimate of the extent to which DRP models can support patient-specific treatment prioritization.

Within this framework, one possible explanation for the strong performance of DWM-NELLY is that the DWM conditions gene contribution to drug chemical features, thereby enabling a drug-specific interpretation of transcriptomic input. This conditional weighting strategy, alongside the compact yet biologically informative representation provided by the L1000 gene set [35], may capture transcriptome-drug interactions more efficiently than heavier architectures, such as those used in attention-based architectures. Supporting this view, a recent benchmark of foundational models for cancer DRP showed that, despite their greater architectural complexity, performance under unseen-CCL evaluation reached an average correlation of only 0.86 [82]. Strikingly, after fixed-drug aggregation, performance decreased remarkably to a correlation of 0.34 [82]. Because the DWM is linearly structured, it also enables direct downstream analysis of DWM-derived gene attribution. This provides a new opportunity to improve model interpretability and support biological investigation at the patient level.

We further used our HNSCC PDO cohort as a case study to explore the biological relevance of DWM-NELLY. In this context, DWM-NELLY prioritized several therapeutic agents with prior biological or clinical relevance in HNSCC, including Pelitinib, Cabozantinib and Tipifarnib [70, 71, 72, 73, 74, 7, 78, 79, 80]. Notably, Tipifarnib has been investigated extensively in HRAS-driven HNSCC [7, 78, 79, 80], supporting the biological plausibility of the predicted recommendations. Beyond drug ranking, analysis of DWM-derived gene attribution suggested at least two modules associated with resistance-related states, including epigenetic/chromatin-regulatory programs and repair- or stress-adaptive programs. These findings were consistent with previously described resistance-associated mechanisms [46, 47, 48, 58, 59, 60, 61, 62, 63, 64] and suggested that DWM-derived gene attribution may provide a useful framework for exploring resistance biology in a patient-specific manner.

Despite these advances, the current study remains limited to preclinical systems rather than to direct prediction of patient response. Although PDOs provide a more clinically relevant model than CCLs, they do not fully recapitulate patient tumors [25, 83, 84]. A major limitation is the incomplete representation of the tumor microenvironment, which is likely to be critical for faithful prediction of treatment response *in vivo* [85, 86]. In addition, the current PDO-based atlas remains limited in its coverage of tumor types and pharmacological space. While the GDSC [31] training resource includes a broad range of disease context, including lung, breast and hematological cancers, the available PDO validation data covered fewer cancer entities. Similarly, the GDSC includes multiple compounds targeting the same pathway, whereas the current PDO atlas provides more limited coverage of therapeutic classes and chemical diversity. This limitation does not primarily concern model training, but rather the range of PDO lines and drug classes available for a clinically relevant assessment of the predictions. Expanding the PDO atlas across additional cancer types, drugs and pathway classes will therefore be important for more comprehensive translational validation of DRP models. Another limitation is that we did the analysis just with tumor organoids and did not assess responses by normal organoids. Combining these into the model could enable the identification of cancer-specific vulnerabilities.

Future work should focus on narrowing the remaining gap between preclinical and patient-derived drug response prediction. In particular, domain adaptation strategies may help improve the transfer of representations learned from CCLs to PDOs and other patient-derived systems. However, the performance difference observed between CCLs and PDOs may also reflect limitations in the current PDO-based pharmacogenomic atlas. In line with this, Xia et al (2022) showed that scarce drug-response measurements can reduce model accuracy and increase predictive uncertainty [21]. Additionally, it will also be important to investigate whether DWM-derived gene attribution can help to identify biomarkers of sensitivity or resistance for specific therapeutic agents. Such candidate biomarkers should then be evaluated experimentally and assessed for their prevalence and relevance across patient populations. Furthermore, Cheng et al. (2019) showed that network-based analysis can be used to predict effective drug combinations. In this context, DWM-derived gene attribution may provide a useful basis for extracting drug- and patient-specific gene fingerprints. These fingerprints could potentially be integrated into network-based frameworks to identify candidate combination therapies in a patient-specific manner [87].

Taken together, our study addresses key limitations in current DRP research for translational applications. By combining patient-centric benchmarking on both CCLs and on PDOs, we demonstrate that DWM-NELLY improves patient-specific drug prioritization across preclinical and patient-derived systems. At the same time, the DWM extends the model beyond prediction by allowing patient-specific biological interpretation. These results support DWM-NELLY as a promising patient-relevant framework for interpretable drug response prioritization in precision oncology.

## Materials and methods

### Patient-derived organoid generation

#### Collection of clinical samples

Resection samples from nine patients with oral squamous cell carcinoma were collected in collaboration with the Department of Oral and Maxillofacial Surgery at the University Clinic Würzburg. Written informed consent was obtained from all patients. The Ethics Committee of the University of Würzburg approved the sample collection as project approval 218/16-me in accordance with national and European legislation. All samples were pseudonymized and the associated clinical data (age, sex, diagnosis and treatment) were correlated. The collected resection samples included treatment-naïve primary tumour samples from various sites in the oral cavity, as well as metastatic samples from cervical lymph nodes. Fresh tissue samples were stored in ice-cold adDMEM+++ (advanced DMEM/F12 (Thermo Fisher Scientific, Cat# 12634-028) supplemented with 10 mmol/l hepes (Thermo Fisher Scientific, Cat# 15630-056), 1 × Glutamax (Thermo Fisher Scientific, Cat# 35050061), 1 × penicillin/streptomycin (Thermo Fisher Scientific, Cat# 15140122)) and processed within 24 h. A representative tissue piece (∼ 1 mm × 1 mm) was cut into smaller pieces. One part was embedded in Tissue-Tek^®^ O.C.T. Compound (Sakura, Cat# 4583), snap-frozen on dry ice, and relocated to −80 *^◦^*C for subsequent histological analysis. The remaining tissue was processed either to isolate epithelial cells and generate patient-derived organoids, or to mince and transfer it into 1 ml of CryoStor^®^ Cell Cryopreservation Media (Sigma-Aldrich, Cat# C2874) for long-term storage at −150 *^◦^*C until further use.

#### Generation of patient-derived oral squamous cell carcinoma organoids

Organoid cultures were established and maintained as previously described by Harnischfeger et al. (2025) [88]. In short, cryopreserved or fresh tissue samples were cut into pieces measuring approximately 0.5–1 mm. These pieces were then enzymatically digested using ad-DMEM/F12+++, which was supplemented with 2 mg/ml collagenase P (Sigma-Aldrich, Cat# 11213865001), 1 mg/ml collagenase D (Sigma Aldrich, Cat# 1088858001), 0.5 mg/ml DNase I (Roche, Cat# 10104159001) and 10 µM Y-27632 (Hölzel Biotech, Cat# M1817). The digestion process took place at 37 *^◦^*C for 30 minutes. To release the epithelial cells, the tissue pieces were resuspended igorously in the digestion solution every 10 minutes. To enhance digestion, the tissue pieces were then digested in a 0.25% trypsin solution (Thermo Fisher Scientific, Cat# 15090046), diluted in phosphate-buffered saline (PBS) (Sigma-Aldrich, Cat# D8537), until they broke down into smaller clumps of 2–10 cells. These were then embedded in 70% Cultrex BME type 2 (bio-techne, Cat# 533-005-02). Once the BME had solidified (∼30 minutes at 37 *^◦^*C), organoid culture medium was added to cover the BME droplets. Organoids in the first five passages were cultured in either OSCC-1 or OSCC-2 medium. The OSCC-1 organoid medium was similar to the medium used by Driehuis et al. (2019) [34], but with some minor adaptations. The OSCC-1 medium contained adDMEM/F12+++, supplemented with 20% R-Spondin 1-conditioned medium (made in-house), 10% Noggin conditioned medium (made in-house), 1x B-27^TM^ supplement (serum-free, Thermo Fisher Scientific, Cat# 17504001), 1.25 mM N-acetyl-L-cysteine (Thermo Fisher Scientific, Cat# A9165), 10 mM nicotinamide (Sigma-Aldrich, Cat# N0636), 50 ng/ml human EGF (PeproTech, Cat# AF-100-15), 500 nM A83-01 (Bio-Techne, Cat# 2939), 10 ng/ml FGF10 (Qkine, Cat# Qk003-1000), 10 ng/ml human FGF2 (PeproTech, Cat# 100-18B), 1 µM prostaglandin E2 (Bio-Techne, Cat# 2296), 3 µM CHIR99021 (Sigma-Aldrich, Cat# SML1046), 1 µM forskolin (Bio-Techne, Cat# 1099), 25 ng/ml human FGF-7 (Thermo Fisher Scientific, Cat# 100-19), 1 µM (SB202190 Sigma-Aldrich, Cat# S7067). The OSCC-2 medium composition resembled the endocervical M7 medium used by Lõhmussaar et al. (2021) [89] and was supplemented with 0.5 nM WNT surrogate (ImmunoPrecise Antibodies, Cat# N001), 50 ng/ml EGF (PeproTech, Cat# AF-100-15) and 0.3 mM CHIR99021 (Sigma Aldrich, Cat# SML1046). Stable, well-expanding organoid lines that had reached passage five or further were subsequently transferred to OncoPro^TM^ Tumoroid Culture Medium (Thermo Fisher Scientific, Cat# A5701201) [90]. o avoid contamination, 100 µg/ml Primocin (InvivoGen, Cat# Ant-pm-1) and 0.5 µg Caspofungin (Sigma Aldrich, Cat# SML0425) were added to all three medium conditions, independently of the organoids’ passage number. Passaging of organoids followed the same procedure as described by Harnischfeger et al. (2025) [88].

### RNA-sequencing

#### RNA isolation and bulk RNA-seq library preparation

The total RNA of the organoids was isolated using the NucleoSpin RNA Kit (Macherey-Nagel) according to the manufacturer’s instructions. The quantity and quality of the RNA was assessed with the NanoDrop One (Thermo Fisher Scientific), the Qubit^TM^ RNA High Sensitivity (HS) Kit (Thermo Fisher Scientific) and the BioAnalyzer (Agilent). Library preparation for bulk RNA sequencing followed the CEL-Seq2 protocol by Hashimshony et al. (2016) [91] with minor adaptions. 8 ng of total RNA was used for sequencing. For library preparation, 10 ng/µl of aRNA was reverse transcribed. Ten PCR cycles were used to amplify the amount of cDNA. The final DNA concentration was measured using the Qubit^TM^ 1X dsDNA High Sensitivity (HS) Kit (Thermo Fisher Scientific) and the BioAnalyzer (Agilent High Sensitivity DNA Kit).

#### Bulk RNA-seq mapping and counting

The methods for this analysis were conducted according to the procedures detailed in the publication by van den Brink et al. (2017)[92]. Initially, raw FASTQ files were concatenated specifing the lengths of the cell barcode (CBC) and the unique molecular identifier (UMI). To ensure high-quality reads, adapter sequences were removed using TrimGalore (Version 0.6.8), which utilizes cutadapt for trimming. Subsequently, a genome index was created using the STAR aligner (Version 2.7.0a), with the human genome assembly GRCh38 and its corresponding GTF file serving as references. The trimmed reads were then mapped to the human genome using STAR. The output was generated in BAM format and sorted by coordinate, including all SAM attributes and intron motifs, and retaining only uniquely mapped reads for further analysis. Exon and intron regions were generated from GTF file to generate as BED. Gene counting and annotation were performed using the aligned BAM files and the BED files generated for exons and introns. This was accomplished with the aid of bedtools, which counted the reads mapping to exon and intron regions, producing the count matrix.

### Datasets

#### Training dataset

For model training, cancer cell line drug response data from the Genomics of Drug Sensitivity in Cancer (GDSC) [93, 31] database were paired with gene expression data from the Cell Model Passports database [94].

Drug response metrics are provided as half-maximal inhibitory concentration (IC_50_) values, one of the most widely used metrics for drug response and prediction [95, 96]. Raw IC_50_ values were natural logarithm (ln) transformed and Z-scored. When CCL-drug pairs appeared in both versions GDCS1 and GDCS2, only the response of the GDSC2 was retained. For each chemical compound in the dataset, its canonical Simplified Molecular Input Line Entry System (SMILES) was obtained by querying PubCHEM [97, 98, 99] using GDSC-provided compound identifiers. The canonical SMILES representation was converted to Morgan fingerprints of 512 bits using RDKit [36, 100]. The final dataset contained 952 different CCLs and 413 different drugs.

Each CCL was represented by an array of expected counts that was normalized to counts per million (CPM) and *log*_2_ transformed (with a pseudocount of 1). Gene expression matrix was then filtered to retain only the landmark genes included in the L1000 platform [35], which were originally selected to provide a compact representation of transcriptional variation across cellular stated and perturbations.

#### Cross-validation

To emulate clinical applications for previously unseen biological models, the dataset was split using an unseen-CCL 10-fold cross-validation fashion. In each cross-validation iteration, CCLs were assigned to three mutually exclusive sets: eight folds where used for training, one fold for validation and one fold for testing, corresponding approximately to an 80-10-10 sample split. Furthermore, splits were stratified by tumor type to ensure unbiased model learning of individual diseases.

#### Independent dataset

Studies describing PDO lines for colorectal cancer [25], bladder cancer [32], pancreatic cancer [33] and head and neck squamous cell carcinoma [34] were mined for drug response and gene expression data to establish an independent benchmark dataset. Raw gene expression count data for colorectal and bladder cancer were downloaded from Gene Expression Omnibus (GEO) using accession numbers GSE64392 and GSE103990, respectively. Raw gene expression count data for pancreatic cancer were provided by the authors of the above-referenced study. For head and neck squamous cell carcinoma, raw gene expression count data were partly provided by the authors of the above-referenced study and, additionally, *de novo* generated in this study following protocols described in Driehuis et al. (2019) [34].

Drug response values were harmonized by ln-transformation and Z-scoring of IC_50_ values. Furthermore, raw gene expression counts were normalized to counts per million and *log*_2_-transformed (with a pseudocount of 1). All preprocessing steps were implemented in Python using NumPy, [101].

### Model architecture

#### Naive-NELLY

NELLY consisted of a multimodal neural network with three dedicated heads: a gene expression (GEX) encoder, a chemical encoder and a predictor head (Fig. 1C). During model development, NELLY was trained on CCL-drug pairs. At inference, the same formulation can be applied more generally to line-drug pairs, where a line corresponds to either a CCL or a PDO line.

The gene expression head *G* : X_GEX_ → R*^d^* encoded the gene expression profile *x_i_*^GEX^ ∈ X_GEX_ of CCL *i* ∈ L_CCL_ into a latent embedding *Z*_GEX_ ∈ R*^d^*. The architecture included 3 fully connected layers with 512, 256, 1024 neurons. Each fully connected layer was followed by Leaky ReLU activation.

The chemical head *S* : X_SMILES_ → R*^d^* encoded the drug chemical profile *x_j_*^SMILES^ ∈ X_SMILES_ of a drug *j* ∈ D into a latent embedding *Z*_SMILES_ ∈ R*^d^*. The architecture included 3 fully connected layers with 512, 256, 128, 1024 neurons. Each fully connected layer was followed by Leaky ReLU activation.

We assume an additive structure of the drug response, as previously discussed by Hetzel et al. (2022)[102], therefore the *Z*_GEX_ and *Z*_SMILES_ embeddings were then summed and fed into the predictor head *H* to predict a Z-score ln(IC_50_) value. The architecture included 2 fully connected layers with 256, 128 neurons. Each fully connected layer was followed by Leaky ReLU activation.

The model was trained for 100 epochs using the mean squared error (MSE) loss function, with a learning rate of 1 × 10*^−^*^4^, a batch size of 512.

#### DWM-NELLY

Additionally, NELLY included a specialized module designed to boost model performance and improve model transparency. The module was referred to as Dynamic Weighting Mechanism (DWM), because it generated gene weights for each line-drug pair, dynamically adjusting gene contribution to each transcriptomic and chemical profile. Unlike attention-based architectures, which typically aggregate information through latent or pathway-level representations, the DWM applies a drug-conditioned gene-wise gate directly to the transcriptomic input (Fig. 2A). This preserves a direct correspondence between each learned weight and an input gene, allowing the resulting weights to be analyzed as pair-specific gene attributions. The DWM can be described mathematically as follows:

Let *X*_GEX_ ∈ R*^N×d^* denote the gene expression matrix and *Z*_SMILES_ ∈ R*^N×dz^* the matrix of drug embeddings. Where *N* denotes the number of CCL-drug pairs. The two representations were concatenated along the feature dimension to form

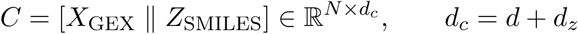

The concatenated representation was then projected through a linear layer followed by a sigmoid function

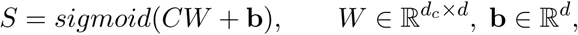

were *W* and *b* denote the learnable parameters. This produced a gene-weight matrix *S* ∈ R*^N×d^*. The original input *X*_GEX_ was rescaled by the gene-weight matrix *S* through element-wise multiplication, and summed back through residual connection [37] for model stability:

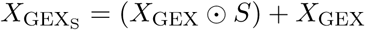

This operation is equivalent to scaling the original input by 1 + *S*, but writing it as a residual connection makes explicit that the original transcriptomic signal was preserved. Therefore, the DWM learned a drug-conditioned gene-attribution matrix (*S*) that modulated each gene’s contribution into the prediction task. Gene weighting served as a gating step applied in a CCL- and drug-specific manner.

#### Benchmarking with existing deep learning models

For benchmarking model performance, we selected three existing deep learning (DL) models for drug response prediction: Paccmann [13, 14], Hierarchical Network for Drug Response Prediction with Attention (HiDRA) [15] and ScreenDL [29]. These models were selected because they were originally developed and evaluated using GDSC drug response data. Additionally, they used gene expression data as the molecular input and integrated drug chemical information, making them directly comparable to NELLY. Together, they represent distinct state-of-the-art strategies for DRP, including multimodal multi-head attention-based integration, pathway-informed hierarchical attention and deep neural drug-response modeling.

To ensure a rigorous comparison, all benchmark models were retrained on the same curated GDSC dataset used for NELLY. For each model, CCL and drug representations were generated according to the preprocessing and feature-construction procedures described in the corresponding publication. Models were retrained using the provided hyperparameters and evaluation was done under the same unseen-CCL 10-fold cross-validation scheme. Therefore, all models were compared using identical CCL-level train-validation-test splits. Pretrained models were not used because differences in training data, preprocessing, and splitting strategy would have confounded architectural comparisons.

Model performance was first evaluated using the Pearson correlation coefficient (PCC) between observed response (*Y*) and predicted (*Ŷ*) response. The PCC is computed as follows:

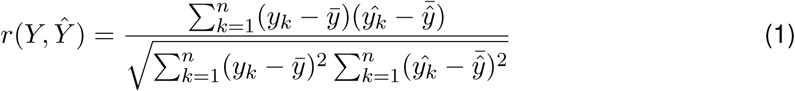

where

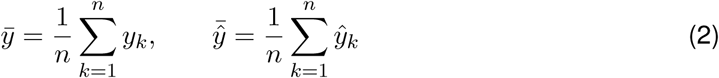

Previous studies have shown that unstratified evaluation can overestimate DRP model performance [19, 16]. We therefore adopted aggregation-based strategies, using *Fixed-drug* and *Fixed-line* schemes as described by Codicè et al.(2025) [16]. Throughout the benchmarking section, we use the term line as a generic term to refer to either biological line, i.e., cancer cell lines in the training benchmark or PDO lines in the independent benchmark.

Let *D* denote the set of drugs *j* ∈ [1*, …, m*] and *L* denote the set of biological lines under evaluation *i* ∈ [1*, …, n*]. Depending on the dataset, lines correspond either to cancer cell lines or organoid lines. Then, for line *i* ∈ *L*, let *D_i_* ⊆ *D* denotes the set of drugs tested in line *i*. For drug *j* ∈ *D*, let *L_j_* ⊆ *L* denotes the set of lines exposed to drug *j*. Then, for a fixed drug *j* ∈ *D*, across lines *i* ∈ *L*, PCC is computed as:

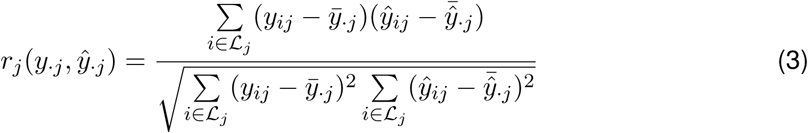

where

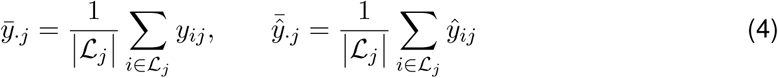

On the other hand, for a fixed line *i* ∈ *L*, across drugs *j* ∈ *D*, PCC is computed as:

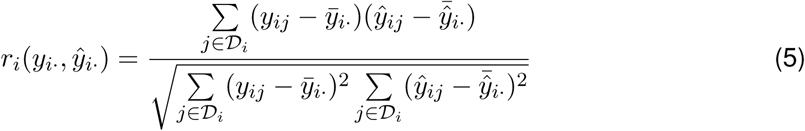

where

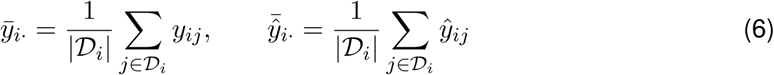

For model comparison across the 10-fold cross-validation scheme, the median PCC value was taken for both aggregation strategy and statistical comparison was done with two-sided Wilcoxon signed-rank test. Therefore:

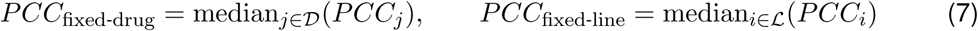

To complement regression accuracy, we evaluated ranking performance using Precision and Normalized Discounted Cumulative Gain (NDCG) [103]. Following the results from Lee et al. (2018), we defined the most sensitive responses as those falling within the lower 25% (Q25) of the Z-score ln(IC_50_) distribution, recognizing that lower response values indicate greater sensitivity [104]. For both Precision and NDCG, we set relevance to the Q25 of the observed response distribution. Therefore, these metrics measure how well a model detects and orders the most sensitive responses. Ranking accuracy was evaluated under both fixed-drug and fixed-line aggregation. Given fixed-drug aggregation, items correspond to the lines exposed to a given drug *L_j_*. Under fixed-line aggregation, items correspond to the drugs tested in a given line *D_i_*. Let *S_u_* denote the candidate set for ranking unit *u*, where *u* = *j* for fixed-drug aggregation and *u* = *i* for fixed-line aggregation.

The relevant set *R_u_* ⊆ *S_u_* was defined as the items whose observed response values fell within the lower 25% of the observed response distribution for unit *u*. Let *k_u_* = |*R_u_*| denote the number of relevant items. Predicted responses were ranked in ascending order, and the top-*k_u_* predicted items were taken as the predicted sensitive set *R̂_u_*.

Precision@Q25 for unit *u* was computed as:

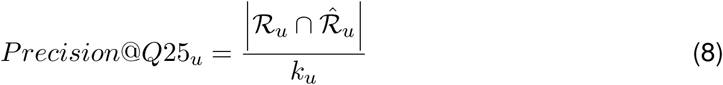

To evaluate the ordering of the predicted sensitive items, we also computed NDCG@Q25. Relevance was defined as the degree of observed sensitivity. Because the response variable corresponded to Z-score ln(IC_50_), lower values indicated stronger sensitivity. Therefore, the observed response values were converted into relevance scores by reversing their sign rel*_u_*(*s*) = −*y_u,s_*. Following this transformation, the more sensitive line-drug pairs received higher relevance values, where *y_u,s_* denotes the observed Z-score ln(IC_50_) value for item *s* in ranking unit *u*. The discounted cumulative gain for unit *u* was then computed over the top-*k_u_* predicted items as:

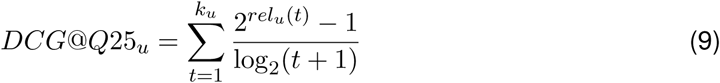

where rel*_u_*(*t*) denotes the observed relevance of the item ranked at position *t* by the model.

The normalized discount cumulative gain was then obtained as:

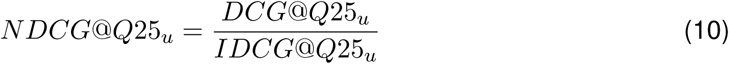

where *IDCG*@*Q*25*_u_*is the ideal discounted cumulative gain obtained by ranking items according to the observed response values.

For model benchmark across the 10-fold cross-validation, median ranking performance was computed similarly to PCC and statistical comparison was done with two-sided Wilcoxon signed-rank test.

## Data and code availability

Training data: drug response data were downloaded from https://www.cancerrxgene.org/, gene expression data were downloaded from https://cellmodelpassports.sanger.ac.uk/. Pan-cancer PDO pharmacogenomic atlas: raw gene expression count data for colorectal and bladder cancer were downloaded from Gene Expression Omnibus (GEO) using accession numbers GSE64392 and GSE103990, respectively. Raw gene expression count data for pancreatic cancer were provided by the authors [33]. For head and neck squamous cell carcinoma, raw gene expression count data were partly provided by the authors [34] and, additionally, *de-novo* generated in this study. Data generated in this study is available upon request to the lead contact. Raw sequencing data is subject to a material transfer agreement due to patient data protection and confidentiality.

All code used for model development and analysis is available at https://github.com/cperalta-viteri/NELLY.

## Acknowledgment

We thank Dominic Grün and Christophe Zimmer for their valuable feedback and critical comments on the manuscript. We thank Emilia Stanojkovska for the hand-drawing illustrations. We thank Mara John, Nazia Afrin, Moutaz Helal, Martin Eilers and Martin Czolbe for their advice and help. We acknowledge technical support from the Core Units for Systems Medicine at the University of Würzburg.

## Funding

This work was supported by funding from the German Cancer Aid (MSNZ Würzburg/NG to K.K) and the European Union/European Research Council (ERC Starting Grant number 101042738/OralNiche to K.K.). C.P.V., N.H. and L.S. were supported by funds from the Bavarian State Ministry of Science and the Arts and the Graduate School of Life Sciences (GSLS) of the University of Würzburg.

## Author contributions

C.P.V.: conceptualization, methodology, software, validation, formal analysis, investigation, data curation, visualization, writing—original draft. N.H.: investigation, data curation, writing—original draft. L.S.: investigation, data curation, writing—review and editing. S.H.: resources, data curation, writing—review and editing. K.K.: conceptualization, supervision, funding acquisition, project administration, writing—original draft. All authors reviewed and approved the final manuscript.

## Notes

### Competing Interest Statement

The authors have declared no competing interest.

